# Preparation of Artesunate mPEG-PLGA Nanoparticles and Its Application to K562 Apoptosis of cells

**DOI:** 10.1101/628883

**Authors:** Sushil Reddy, David Gupta

## Abstract

The preparation process of artesunate-loaded polyethylene glycol monomethyl ether-polylactic acid-glycolic acid affinity block copolymer (mPEG-PLGA) nanoparticles and its growth inhibition on human leukemia K562 cells were investigated. METHODS: Artesunate mPEG-PLGA nanoparticles (Art-Nps) were prepared by modified self-emulsification method. The morphology of nanoparticles was characterized by scanning electron microscopy. The particle size distribution and zeta potential were measured by laser scattering particle size analyzer. The drug loading, encapsulation efficiency and in vitro release of Art-Nps were determined by chromatography. The proliferation and apoptosis of human leukemia K562 cells were observed by MTT assay and Hoechst staining. RESULTS: Art-Nps is a spherical solid particle with smooth surface, average particle size (156.70+/-1.01) nm, zeta potential of -(26.23+/-1.86) mV, average drug loading (14.51+/-0.20)%, average package. The sealing rate was (86.51+/-0.50)%, and the in vitro release law accorded with the Higuchi equation: Q=4.11t 1/2+27.05, R2=0.98. MTT assay showed that Art-Nps inhibited the proliferation of K562 cells in a time-dose-dependent manner, and the inhibition rate exceeded the artesunate-treated group after 72h, and sustained release. The number of cells was observed after cultured with different concentrations of Art-Nps for 48h. Significantly reduced, cell size is different, irregular shape, high magnification can be seen in the nucleus pyknosis, agglutination, and apoptotic bodies, and increased apoptotic bodies with increasing concentration.

## Introduction

Artesunate (Art) is an artemisinin-based antimalarial drug with self-developed sesquiterpene structure. In recent years, it has been found to have a variety of tumors such as leukemia, colorectal cancer and oral cancer. Strong inhibition, especially for leukemia cells [1], has been included in the anticancer drug screening and anticancer activity research program by the National Cancer Institute. However, Art can cause peripheral blood reticulocyte reduction and teratogenic and other toxic side effects, and the half-life is only 30min, the bioavailability is low, in order to reduce the toxicity of the drug, prolong the half-life of the drug, improve the bioavailability, and reduce the number of administrations. It is made into biodegradable nanoparticles (nanopaticles, Nps). The biocompatible and biodegradable polyethylene glycol monomethyl ether-polylactic acid-glycolic acid affinity block copolymer (mPEG-PLGA) was used as the carrier to prepare Art-mPEG-PLGA nanometer by self-emulsification method. Granules, physicochemical properties were studied, and artesunate was used as a control to study the growth inhibition and apoptosis-promoting effect of drug-loaded nanoparticle solvent on leukemia K562 cells, aiming to explore the anti-leukemia effect of new artesunate dosage form. The value on.

## Materials and Method

Artesunate (Guangxi Guilin Nanfang Co., Ltd., batch number: 091129-6); polyethylene glycol monomethyl ether-polylactic acid-glycolic acid block copolymer (mPEG-PLGA, mPEG molecular weight 5 000, 4%), PEG/PLGA=50/50, PLGA molecular weight 15 000, Shandong Jinan Biotechnology Co., Ltd.); Artesunate reference product (China Biological Products Inspection Institute); Poloxamer F-68, thiazolyl (MTT)), dimethyl sulfoxide (DMSO) (all Sigma, USA); dialysis membrane (molecular weight cutoff 14 000); calf serum (Hangzhou Sijiqing); PRM-1640 medium (Gibco, USA); other reagents All are analytically pure.

Rotavapor R-114 Rotary Evaporator (BUCHI, Switzerland); CL-2 Magnetic Stirrer (Henan Gongyi Yuhua Instrument Co., Ltd.); Zeta-sizer 3 000HS Laser Scattering Particle Size Analyzer (Malvin, UK); TMP Electronics Balance (Satorius, Germany); Centrifuge (THERMO LEGEND, USA); x w- 8 0A vortex oscillator (Jiangsu Haimen Linbei Instrument Co., Ltd.); JSM-6330F scanning electron microscope (Japan Electronics Co., Ltd.); WATERS 600E −2487 High Performance Liquid Chromatograph (Waters Corporation, USA); Freeze Dryer (USA LABCONCO FreeZone 12L); Dissolution Apparatus (HTY-EU802, Germany).

Rotavapor R-114 Rotary Evaporator (BUCHI, Switzerland); CL-2 Magnetic Stirrer (Henan Gongyi Yuhua Instrument Co., Ltd.); Zeta-sizer 3 000HS Laser Scattering Particle Size Analyzer (Malvin, UK); TMP Electronics Balance (Satorius, Germany); Centrifuge (THERMO LEGEND, USA); x w- 8 0A vortex oscillator (Jiangsu Haimen Linbei Instrument Co., Ltd.); JSM-6330F scanning electron microscope (Japan Electronics Co., Ltd.); WATERS 600E −2487 High Performance Liquid Chromatograph (Waters Corporation, USA); Freeze Dryer (USA LABCONCO FreeZone 12L); Dissolution Apparatus (HTY-EU802, Germany).

Preparation of artesunate mPEG-PLGA nanoparticles [2] Artesunate mPEG-PLGA nanoparticles were prepared by self-emulsification method. 10 mg of artesunate powder and 50 mg of mPEG-PLGA were dissolved in 850 μL of acetone and 150 μL of absolute ethanol. After complete dissolution, the oil phase was slowly added dropwise to 10 mL of 0.1% poloxamer in medium-speed magnetic stirring. The mixture was reacted at room temperature for 5 h in an aqueous solution of F-68, and the solution was further evaporated under reduced pressure until the organic solvent was completely evaporated. Finally, a 0.22 μm filter was filtered to obtain a nanocolloid solution. The solution was ultracentrifuged (20 000 r / min) for 0.5 h at 4 ° C, and the precipitate was washed with distilled water, and washed three times with the same method, and lyophilized to obtain artesunate mPEG-PLGA nanoparticles.

Nanoparticle size, span measurement and morphological observation A small amount of washed artesunate nanoparticles were obtained, and the diluted solution was light blue emulsion. The particle size distribution and zeta potential of the nanoparticles were measured using a laser scattering particle size analyzer. The morphology of the nanoparticles was observed under a scanning electron microscope.

3. Determination of drug loading and encapsulation efficiency of nanoparticles was determined by high performance liquid chromatography (HPLC). The chromatographic conditions were as follows: Column: ECOSIL C18 column (150 mm × 4.6 mm, 5 μm); mobile phase: acetonitrile: 0.02 mol / L ammonium sulfate solution: 12% triethylamine solution (60: 40: 0.2, pH adjusted with dilute ammonia) Value to 4.5); Flow rate: 1.0 mL/min; Column temperature: room temperature; Detection wavelength: 210 nm

## Results and Discussion

Morphology and particle size of artesunate nanoparticles The artesunate mPEG-PLGA nanoparticle colloidal solution has a pale blue opalescence appearance. The results of laser light scattering measurements showed that the surface of the nanoparticles was negatively charged and the Zeta potential was -(26.23±1.86) mV. The particle size distribution is narrow and basically normal distribution, the average particle size is (156.70±1.01) nm, and the multi-distribution index is 0.08 on average, as shown in Fig. 1. Scanning electron microscopy showed that the nanometers were spherical, indicating smoothness, uniform size, and almost no adhesion between the particles, see Figure 2.

**Figure 1.**
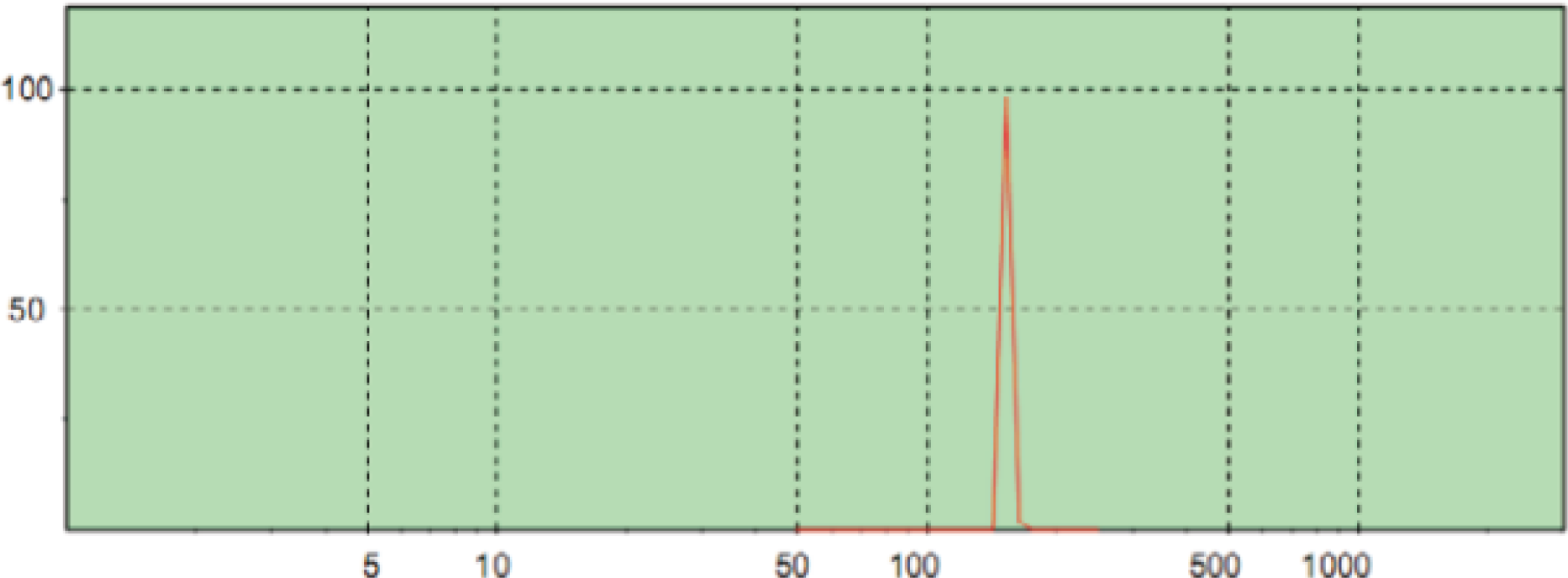
Size analysis by DLS

**Figure 2.**
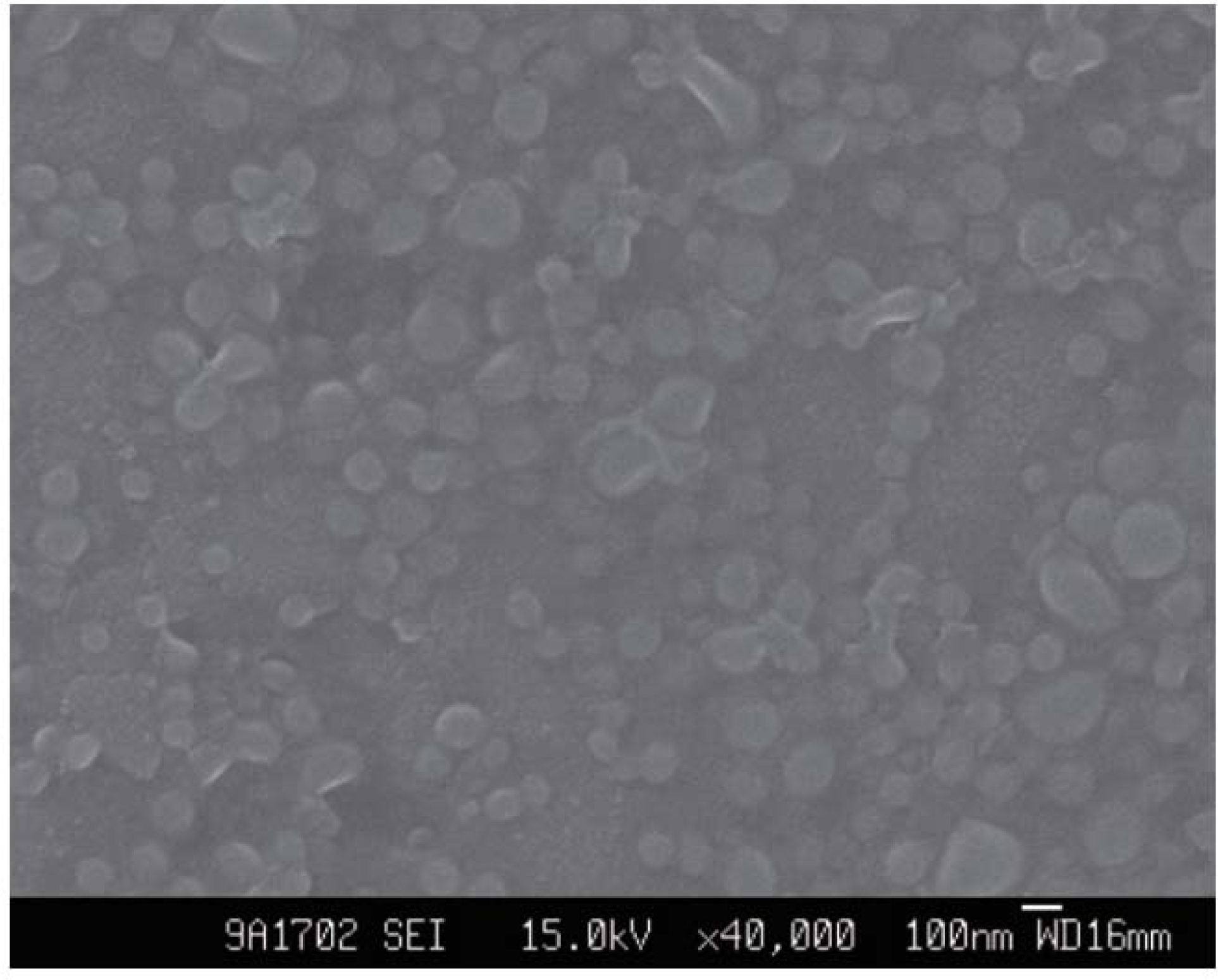
Nanoparticles analyzed by SEM

The drug loading and encapsulation efficiency of artesunate nanoparticles were repeated to prepare 3 batches of artesunate nanoparticles. The average drug loading was (14.51±0.20)%, and the average encapsulation efficiency was (86.51±0.50)%., reproducible.

In vitro release of artesunate nanoparticles in vitro cumulative release curve of artesunate mPEGPLGA nanoparticles, see Figure 3. The simulated internal environment has a burst phenomenon at the beginning of the environment, and the release rate reaches 70% after 5 days of release. By fitting the cumulative release rate and time, the in vitro release of nanometers conformed to the Higuchi equation: Q=4.11t 1/2+27.05, R2=0.98. According to this, the half-life of the release of Art-Nps was 22.61h. The Art-Nps in vitro cumulative release profile is shown in Figure 3. After simulating the in vivo environment for 24 hours, the cumulative release was 45%. After 50 hours, it entered a relatively gentle release process, and the release rate reached 70% after 5 days of release^1-15^.

**Figure 3.**
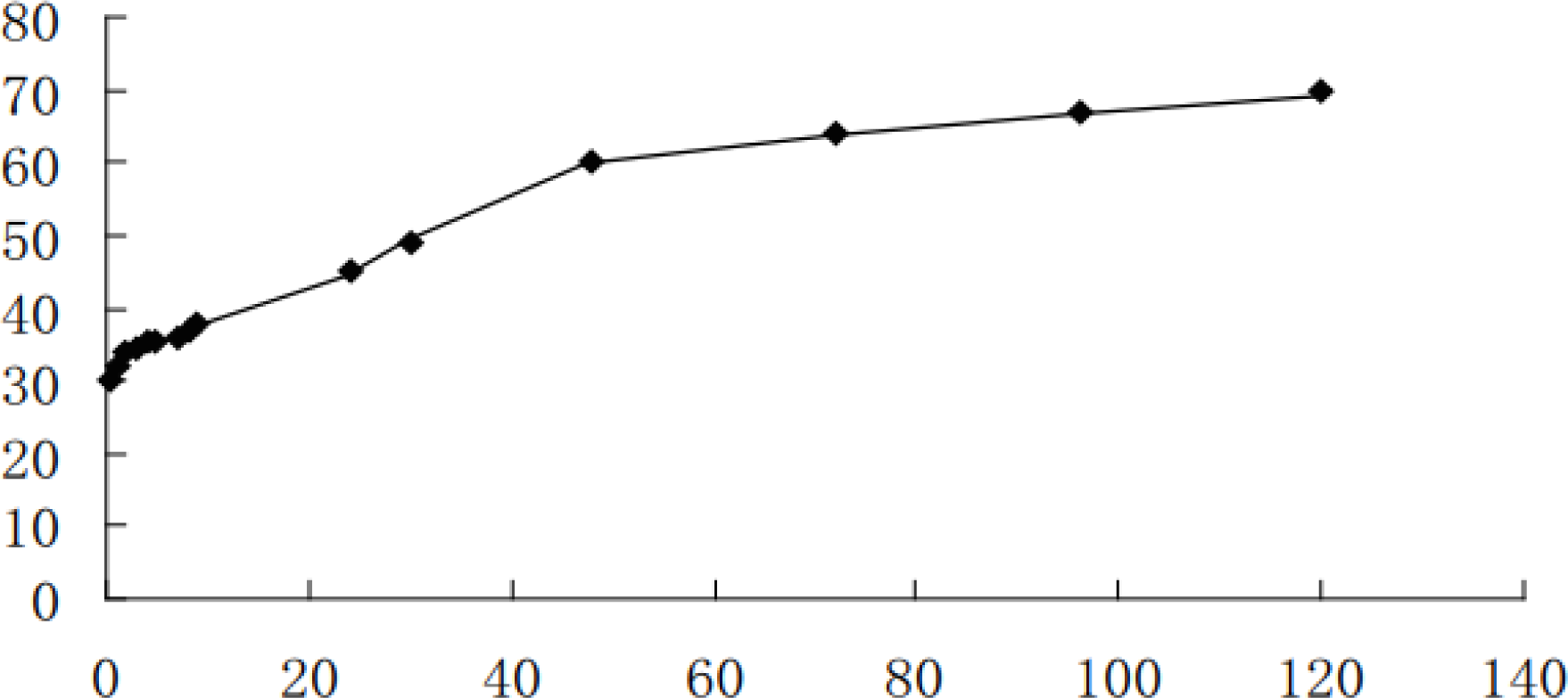
Releasing kinetic analysis

The effect on the growth of K562 leukemia cells compared with different drugs is shown in Figure 4, Table 1. After different concentrations of artesunate nanoparticles were incubated with K562 cells, cell growth was inhibited and showed a dose-effect relationship. Repeated measures analysis of variance and one-way ANOVA showed that different doses (F=407.47, P=0.00) and different drug action times (F=1 301.00, P=0.00) had significant differences in growth inhibition of K562 cells, and There was a significant interaction between the two (F=6.93, P=0.00), suggesting that Art-Nps inhibited the growth of K562 cells in a time-dose dependent manner. Further LSD multiple comparison analysis showed that the Art-Nps concentration treatment group (except for the 50 μg/mL group compared with the 100 μg/mL group) had significant differences in K562 cell inhibition rate at all observation times.

**Figure 4.**
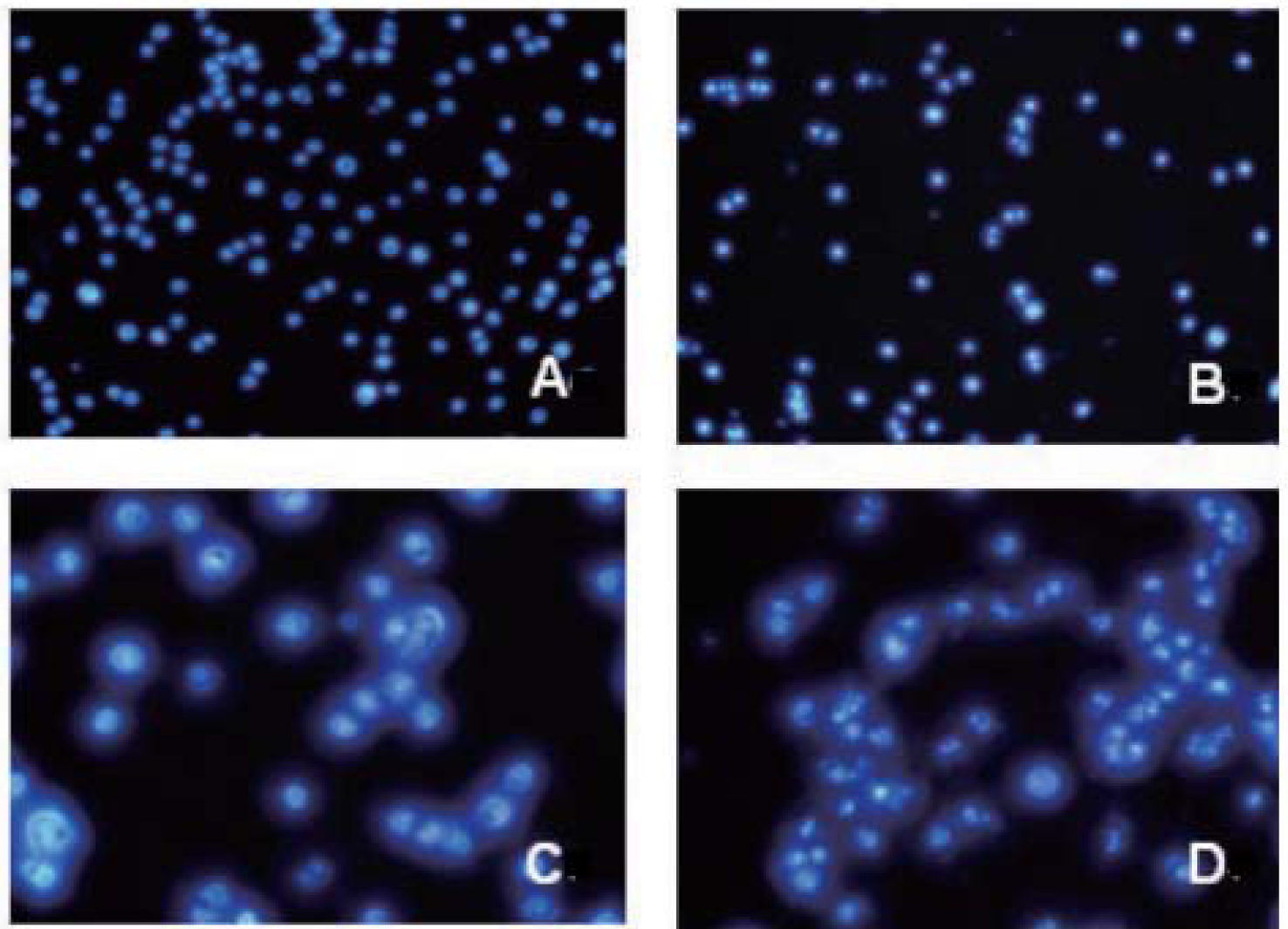
Cell uptake assessed by fluorescent microscopy

Hoechst staining observed changes in the morphology of K562 cells after Art-Nps treatment. In the blank control group, the cells grew well and the number increased with the growth time. The cells were uniform in size, uniform in shape and round in shape. Under high magnification, the nucleus showed diffuse uniform blue fluorescence. After cultured K562 cells with different concentrations (12.5, 25, 50μg/mL) Art-Nps for 48h, the number of cells was significantly reduced, the cell size was different, and the morphology was irregular. The high-power microscope showed nucleus pyknosis, agglutination, and small apoptosis. The body is characterized by bright granular blue staining, and the apoptotic bodies gradually increase with increasing concentration.

The development of nanotechnology has provided a new drug-loading system for biomedicine, which has become a new research hotspot in the field of medicine. At present, there are many researches on nanoparticles loaded with anti-tumor drugs at home and abroad. Anti-tumor drugs include 5-fluorouracil and Azomycin., arsenic trioxide, paclitaxel, etc., and achieved certain results. Artesunate is a research hotspot in recent years as a new type of anticancer drug. Zhang ZY et al [3] reported that artesunate combined with chemotherapy can improve the recent disease control rate and delay disease progression in patients with advanced non-small cell lung cancer. Safety is good; in addition, Sieber S et al [4] reported that artesunate can be combined with meloxicam for non-Hodgkin’s lymphoma, and the two have synergistic effects, but the blood concentration drops rapidly after intravenous injection of Art, half-life is 30min Left and right, repeated administration, so how to develop artesunate sustained-release nanoparticles with anti-tumor activity has become the focus of research.

PLGA and mPEG are two biodegradable biomaterials that can be used in the human body by the US Food and Drug Administration. They are currently used in the preparation of drug-loaded nanoparticles. The two constitute mPEG-PLGA parents through the action of covalent or non-covalent bonds. The block copolymer can effectively improve the hydrophilicity and degradability of the nanoparticles and prolong the circulation time of the nanoparticles in the body [5]. The mPEG is highly hydrophilic, and the content of the copolymer is 2%-5%. Resisting the critical value of protein adsorption, it can produce effective steric stabilization, forming one or more layers of protective hydrophilic coating film, preventing the adsorption of opsonized protein, thereby reducing the purpose of phagocytosis [6], hydrophilic polymerization The molecular chain of the object can also swing freely, forming a protective layer similar to an electron cloud on the surface of the nanoparticle, which better prevents the adsorption of the nanoparticle by the conditioning protein. The PLGA carrier used in this experiment is bound with 4% PEG, which can effectively reduce the phagocytosis of phagocytic cells [7]. It has targeted and sustained release properties, and the immunogenicity and antigenicity of PEG are extremely low. Therefore, PEG modification is used. Nano-surfaces will give better results, which is also well validated in this experiment.

The results of this experiment showed that the in vitro killing ability of artesunate and drug-loaded nanoparticles on human leukemia cell line K562 was time- and dose-dependent. At all observation times, the cell inhibition rate results suggest that the larger the Art-Nps dose and the longer the duration of action, the more obvious inhibition of K562 cell proliferation by Art-Nps. The endocytic properties of the nanoparticles and their distribution in the cells are related to the physicochemical properties of the nanoparticles (eg particle size, charge and hydrophilicity), concentration, incubation time, temperature and cell line. In general, the smaller the particle size, the easier it is to be swallowed by the cells, and the greater the negative charge, the more likely it is to be captured by cells with positive charges on the surface. Usually increasing the concentration of nanoparticles can increase the amount of nanoparticles that are endocytosed. Before reaching saturation concentration, the concentration of nanoparticles is doubled, and the amount of cells that are endocytosed is doubled, but as the concentration of particles increases, endocytosis Efficiency will also gradually decrease. There was no statistically significant difference in the inhibition rate of Art-Nps concentrations from 50 μg/mL to 100 μg/mL in this study, which may be related to the decreased efficiency of cell endocytosis^4-6, 16-30^.

As a sustained release agent, Art-mPEG-PLGA-NP gradually increased its drug release from 1-5d. It was found to be a turning point around 50h, and the release effect was faster before 50h. After 50h, it entered a relatively gentle time. The release process, which is also consistent with the experimental results of the nanoparticles in this paper after 72h inhibition rate over the control group. The experimental results suggest that Art-mPEG-PLGA-NP prolongs the effective time of artesunate on tumor cells by prolonging the release of artesunate over time and maintaining it at a higher concentration. The principle is that in the initial release of nanoparticles in vitro, the drug is mainly released under the concentration gradient. As the nanometer degrades and ablate, the drug diffuses from the pores to the tissue. As the degradation progresses, the nanometer completely disintegrates and the drug is completely Released. The nanoparticles prepared in this experiment showed significant bursts 24 hours before, and the cumulative release percentage was 45%, which may be due to the rapid degradation of the drug adsorbed on the surface of the nanoparticles or the small particle size of the nanoparticle [8].

In order to achieve better therapeutic effect, reduce toxic side effects and prolong the action time of the drug in vivo, this study is the first attempt to make artesunate into drug-loaded nanoparticles with degradable biopolymer and to explore its effect on inhibiting the growth of leukemia cells. And achieved good experimental results. The results show that Art-mPEG-PLGA-NP can be used as a sustained release agent to maintain effective tumor suppressor concentration for a long time without affecting the biological activity of Art, thus anti-tumor for Art-mPEG-PLGANP. Targeted therapy provides the basis.

## Conclusion

The Art-Nps made by this study has the characteristics of small particle size, high drug loading and encapsulation efficiency. It has been shown to induce apoptosis of human leukemia K562 cells in vitro, and it has a sustained release effect and prolongs the drug against leukemia cells. The time of action, this study provides an experimental basis for the development of new formulations of artesunate.

